# Demixing of simultaneously co-expressed four phase-separating proteins in the endoplasmic reticulum lumen

**DOI:** 10.1101/2025.02.03.636337

**Authors:** Haruki Hasegawa

**Affiliations:** Discovery Protein Science, Department of Large Molecule Discovery and Research Data Science, Amgen Inc., South San Francisco, CA 94080, USA

**Author notes:** Corresponding author: Haruki Hasegawa, Ph.D., Amgen Inc., 750 Gateway Blvd, South San Francisco, CA 94080.

**Keywords:** endoplasmic reticulum, protein phase separation, intracellular protein crystallization, protein droplet, protein crystals, inclusion body, neuraminidase-1, cathepsin B, fusolin

## Abstract

Intracellular protein crystallization represents an intriguing form of biomolecular self-assembly. While the list of such crystallizing proteins is growing and their physiological roles are beginning to be elucidated, the underlying requirements and processes for intracellular protein crystallogenesis remain largely unknown. This study examines how simultaneously co-expressed phase-separating proteins influence each other’s phase-separation event in the endoplasmic reticulum (ER) lumen by using four cargoes selected based on their ability to produce distinctive crystals and droplet inclusion bodies. The co-expressed model proteins independently reached their respective threshold concentrations and spontaneously phase-separated into inclusions in the ER without losing their signature morphologic characteristics. The fact that protein crystals and droplets continued to grow in size over time suggests that nascent cargo proteins were continuously synthesized and folded in the ER to fuel the growth of corresponding inclusion bodies. Namely, despite the highly crowded molecular environment, overexpressed cargoes find their mates by self-association and propagate into microscale structures in the ER. This study demonstrates that cells can accommodate up to four distinct phase separation events simultaneously in the ER lumen, and the phase separation events proceed without interfering with each other and without morphological or content mixing.

## 1. Introduction

Intracellular protein crystallization is an intriguing biomolecular self-assembly process. The earliest tractable report was published in 1853 by Charcot and Robin [1] and then in 1872 by Leyden [2] for bipyramidal protein crystals detected in basophils and eosinophils (now called Charcot-Leyden crystals, or CLCs). Their finding was followed by Reinke’s 1896 report on the hexagonal prism-shaped crystals in testicular Leydig cell cancers [3] and by Glaus in 1917 on the immunoglobulin crystals found in multiple myeloma cells [4]. Since then, intracellular protein crystals have been described in many branches of life [5-7] and occurs in humans under both physiological and pathological settings [5, 8].

Compared to the process of membrane-less organelle formation by liquid-liquid phase separation (LLPS) [9, 10], the underlying requirements for intracellular protein crystallization are still poorly understood [11-17]. However, as illustrated below, functional and physiological importance is beginning to be elucidated for various intracellular protein crystals across different cell types and organisms. Firstly, because of dense packing and resistance to proteolysis, the crystalline state is often employed as a space-efficient way to sequester functional proteins at high concentrations. This scheme is employed by insulins in pancreatic β-cells [18], eosinophil’s major basic protein-1 [19], lipoprotein crystals of yolk platelets in amphibian eggs [20], the ER-derived nutritional protein bodies of plants [21], and the insecticidal toxin crystals of *Bacillus thuringiensis* [22]. Secondly, intracellular protein crystals can function as solid-state catalysts. Three examples have been reported, and all three are peroxisome-localizing enzymes. The first is alcohol oxidase from methylotrophic yeast [23]. The second is urate oxidase from rat liver cells [24], and the third is catalase from sunflower cotyledon and potato tubers [25]. Thirdly, intracellular and in vivo protein crystals contribute to pathogenesis and are important in diagnosing diseases such as (1) multiple myeloma, plasmacytoma, myeloma, and lymphocytic leukemia [8], (2) crystal-storing histiocytosis [26], (3) cataracts [27], (4) hemoglobin C disease [28], (5) CLCs that induce type 2 immunity pathogenesis including asthma and chronic rhinosinusitis with nasal polyps [29, 30], (6) neuromuscular diseases such as mitochondrial myopathies [31], (7) Reinke crystals found in Leydig cell tumors [32]. Fourthly, intracellular protein crystals are adapted to fulfill the functional needs of diverse organisms in their unique living environments. For example, several terrestrial and marine bioluminescent organisms use protein crystallization as a shared approach to store enzymes in their light-emitting organs (see [17] and references therein). The fruit fly *Drosophila* uses specialized “crystal cells” that house protein crystals made of Prophenoloxidase 2 [33] for oxygen sensing, storage, and transport [33]. In filamentous fungi *Neurospora crassa*, cells house unique organelles called Woronin bodies composed of HEX-1 protein [34] that seals the plasma membrane when cells are physically damaged [34, 35]. The ciliated protozoan *Paramecium* stores crystalline secretory products named trichocysts that are discharged as a defense mechanism [36]. These examples illustrate that natural selection exploits various strategies to enable protein crystallization inside the cells when advantageous for the cells or the organismal survival. Likewise, in certain disease settings, intracellular protein crystallization is induced due to pathogenesis or causes deleterious symptoms. However, the degree of selection pressure favoring or disfavoring such protein crystallization propensity in wildtype sequences and disease alleles varies considerably [37].

Although the crystallization of one specific protein alone seems challenging enough, cells possess the capacity to support the crystallization of two or more cargoes inside the cells both in recombinant [15, 16] and natural settings [38]. Yet, the cell’s biosynthetic capacity and physical limits have not been fully explored to address how many different protein cargoes can spontaneously phase-separate into crystals or droplets in a selected organelle. To study how cells manage to sort out the mixture of spontaneously phase-separating cargoes in a given subcellular compartment, four biologically unrelated model proteins were selected and co-expressed in different combinations using HEK293 cells. A key experimental design requirement is to select model proteins that produce completely different inclusion body morphology in the ER lumen. Thanks to the stark difference in inclusion body characteristics, it was clear that co-expressed four proteins simultaneously yet independently phase-separated into respective inclusion bodies in the ER without being merged, mashed up, or interfering with one another. The result asserts that proteins can phase-separate into crystals and droplets despite abundant by-stander proteins and other over-expressed cargoes by finding and associating with their mates without content mixing.

## 2. Results

### 2.1. Selection of four model cargoes that spontaneously phase-separate into readily distinguishable inclusion bodies in the ER lumen

Transiently transfected HEK293 cells were shown to possess enough biosynthetic capacity to support at least four independent intracellular protein crystallization events in different subcellular compartments of a single cell [15]. In this study, the same recombinant expression platform was used to extend the previous findings and test whether four different proteins can phase-separate simultaneously in a single subcellular compartment of the cell. Four model cargo proteins were selected based on the following three criteria. (*i*) Each model protein is already known to phase-separate into a distinguishable inclusion body in the ER lumen, and no extensive validation is required anew. (*ii*) Each cargo protein generates inclusion bodies of consistent characteristics unique to the cargo protein, so the given protein phase separation event is readily identifiable based on the inclusion body properties. (*iii*) The inclusion body characteristics of the four model proteins are different enough from each other, so the inclusion body shape or type can be used as a reliable reporter for corresponding phase separation events.

The following four proteins satisfied the above-mentioned criteria and were selected as the model cargoes for this study. (*i*) Human NEU1 that reproducibly generates cubic and square plate-like protein crystals in the ER lumen in various mammalian cells, including HEK293 cells [15, 39] (see Fig. 1A). (*ii*) *Trypanosoma brucei* cathepsin B (Cat B) that crystallizes into square rod-shaped inclusion bodies when overexpressed in Sf9 insect cells [40, 41] and in the ER lumen of HEK293 cells (see Fig. 1B). (*iii*) An insecticidal protein fusolin from *Anomala cuprea* entomopoxvirus (ACEV) that produces spindle-shaped crystals in the ER lumen (Fig. 1C and [16]). (*iv*) An IgM-like multivalent secretory protein, abbreviated as scFv-Fc-stp (N>A) [13], that produces a cluster of spherical protein droplets via LLPS in the ER (Fig. 1D). As shown by the immunofluorescent micrographs in Fig. 1 A-D, all four types of inclusion bodies were encapsulated in the calnexin-positive ER membranes, demonstrating the site of their phase separation is in the ER lumen.

**Figure 1.**
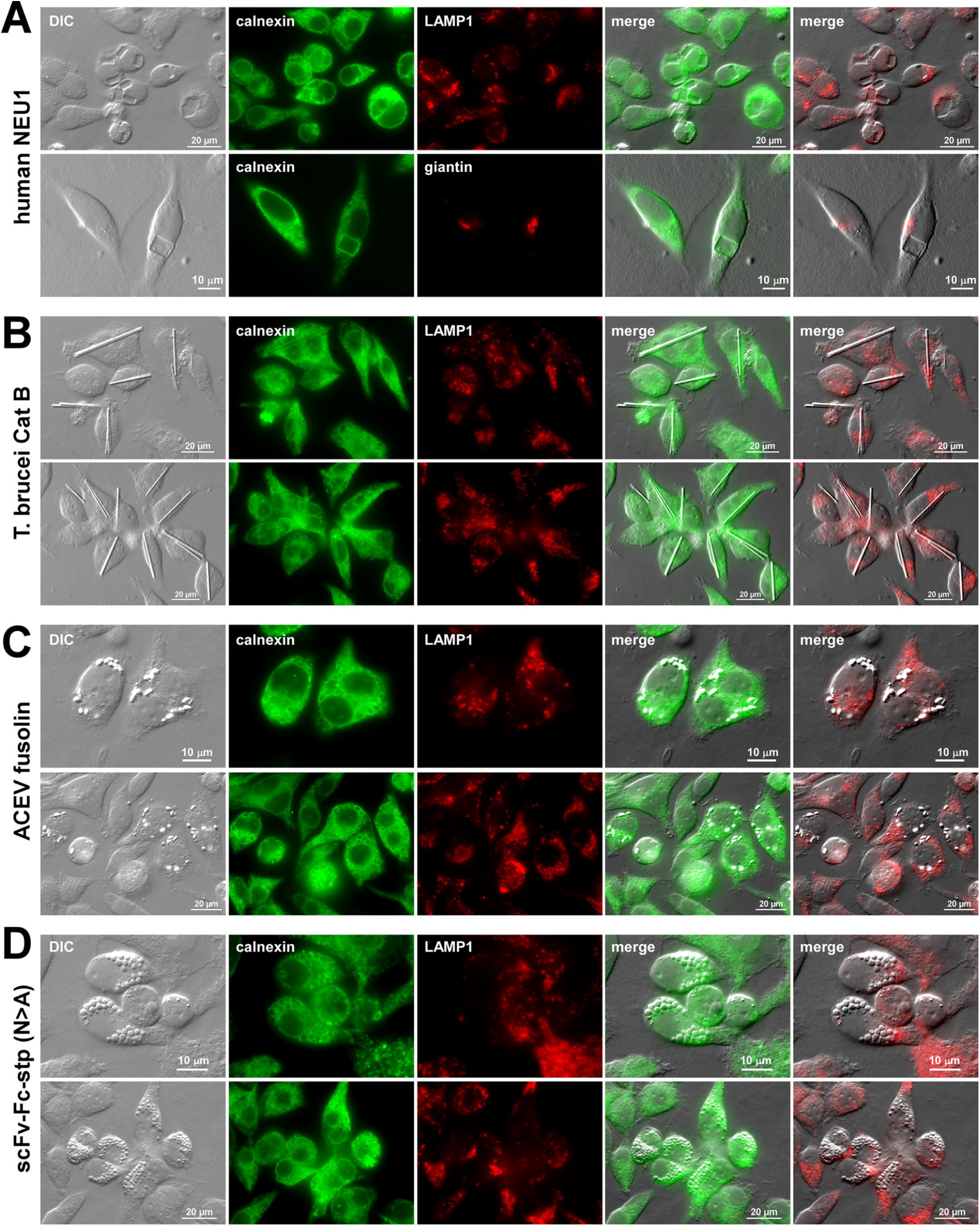
The selected four model cargo proteins produce characteristic and distinguishable inclusion body types in the ER lumen. DIC and fluorescent micrographs of HEK293 cells transfected to overexpress (A) human NEU1, (B) T. brucei Cathepsin B (Cat B), (C) ACEV fusolin, and (D) IgM-like oligomeric secretory protein, scFv-Fc-stp (N>A). On day-3 post-transfection, cells were fixed, permeabilized, and immunostained with rabbit anti-calnexin (shown in green). Co-staining was performed with mouse anti-giantin (shown in red, A, second row) or mouse anti-LAMP1 (shown in red, all the rest). Both green and red images were digitally overlayed with DIC to create ‘merge’ views.

### 2.2. The co-expression of two cargo protein pairs in all six combinations demonstrates there is no interference between any pairs of cargo proteins

The selected four model proteins were co-expressed in pairs in six combinations. This two-gene co-expression study intends to validate that none of the four model cargo proteins prevent other proteins from reaching their respective threshold concentrations in the ER lumen. Similarly, this co-expression experiment preemptively ensures that no two protein cargo pairs are indistinguishable when two morphological types of inclusion bodies are produced side by side in the same continuous ER lumen.

The two-product co-expression experiments demonstrated that individual inclusion body morphology native to each cargo protein was maintained even when co-expressed with another cargo protein in the same ER lumen (Fig. 2). Although a thinning of the individual Cat B rod was observed when it was co-expressed with NEU1 (Fig. 2, top row), there was no other change in inclusion body morphology. Therefore, no two phase-separation events were mixed or mashed into generating ambiguous intermediate inclusion types. Even when two similar inclusion bodies — *i*.*e*., scFv-Fc-stp droplets and fusolin spindles — were formed side by side, they were readily distinguishable based on their appearances (Fig. 2, bottom row). Overall, there were no instances where one protein crystallization event prevented the phase separation of the co-expressed other protein in the ER. In other words, respective cargo proteins could find and associate with their own “mates” in the crowded environment of the ER and produce characteristic inclusion bodies unique to each protein without interfering with one another.

**Figure 2.**
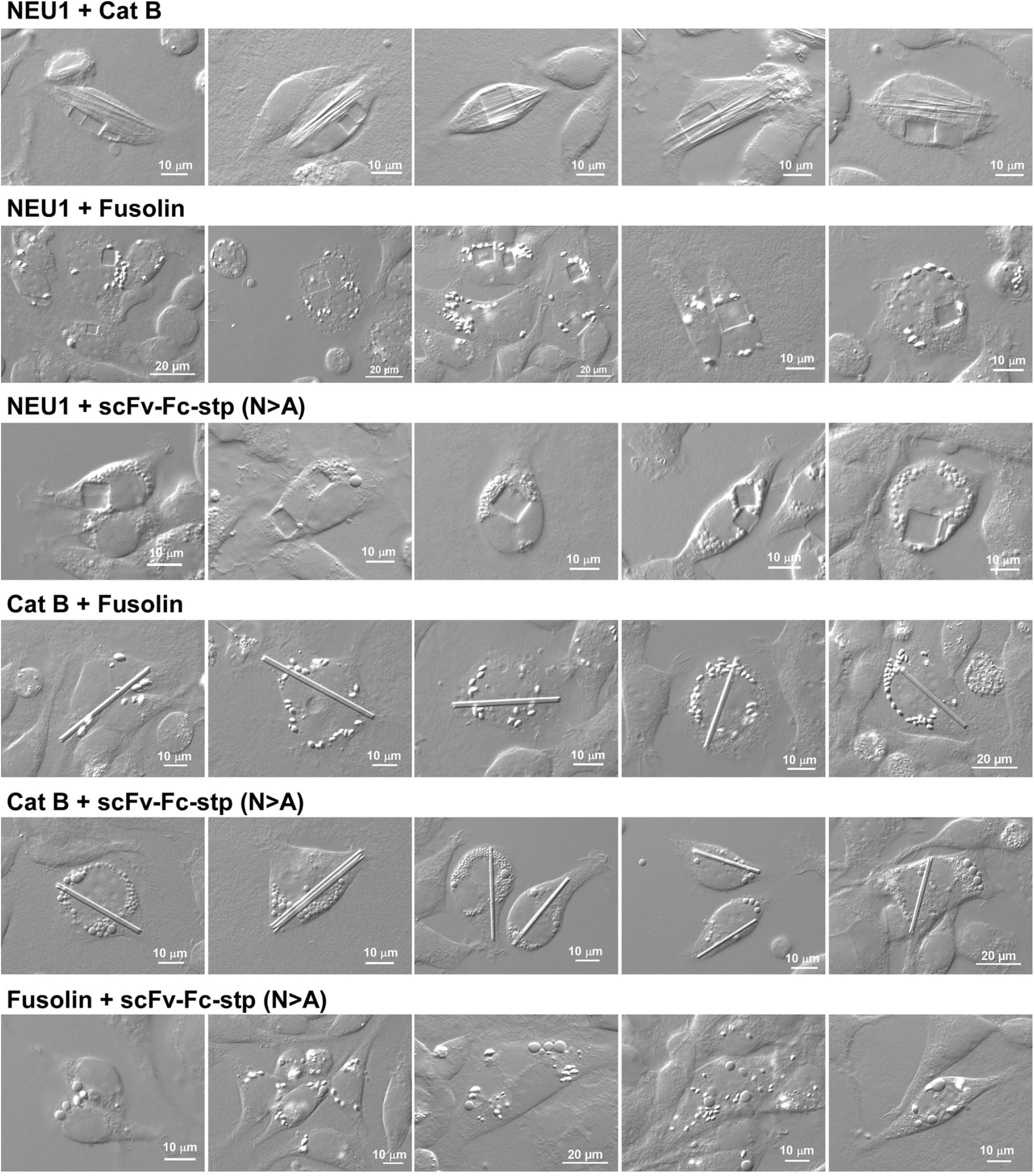
Cargo protein maintains the characteristic inclusion body morphology regardless of the co-expressed cargo proteins. DIC micrographs of transfected HEK293 cells co-expressing the two model cargo pairs in six combinations. On day-4 post-transfection, cells were fixed and imaged. Five representative image fields with the cells of interest are shown for each co-expression pair. Co-expressed protein names are shown on the left side of each row.

Furthermore, these results indirectly suggested that no pairs of inclusion bodies triggered acute ER stress, which otherwise activates the PERK signaling cascade that downregulates cellular protein synthesis [42]. If any of the cargo pairs activated PERK, neither protein pair would have accumulated to a saturating high protein concentration in the ER to induce phase separation events and/or maintain cell viability during the duration of cell culture experiments [43].

### 2.3. The co-expression of all four cargo proteins led to simultaneous phase separation events that do not perturb each other’s inclusion body formation

One of the main purposes of this study is to test whether up to four different types of inclusion bodies can be induced and accommodated simultaneously in the ER lumen of a single cell by challenging the cellular capacity. As noted above, the clue to identifying each phase separation event in simultaneously happening multiple events is assessing the morphology of the inclusion body that reports the protein identity.

When four expression constructs were co-transfected, four different inclusion body types were spontaneously produced in a number of transfected cells by day-4 post-transfection. Although the formation of four distinct inclusion bodies was readily detectable inside the cells frequently, micrographic documentation was not easy because different inclusion bodies were often present in different focal planes of a given image field and usually positioned in various orientations within a cell. As such, determining the precise number of 4-crystal-housing cells was a challenge, and thus, explicitly deriving the percentage of transfected cells that housed 4 different inclusion types was not feasible. Nevertheless, by screening enough image fields, we could document numerous cells that house four distinct inclusion bodies within a depth of two adjacent focal planes apart.

In Fig. 3A, images taken from two adjacent focal planes are displayed side by side for eight different image fields containing the cells of interest that simultaneously house all four inclusion body types. Representative inclusion bodies of the four cargo proteins are pointed out in the image panels. The panels in Fig. 3B show sixteen independent image fields showing the cells of interest that house all four inclusion body types simultaneously in a single focal plane. Representative inclusion bodies for the four cargo proteins are labeled in the two images at the top row. Evidently, each protein reached its threshold concentration in the ER lumen, and hence, four independent phase separation events were simultaneously induced in the ER of a single cell without impacting the signature characteristics of the inclusion body unique to each protein cargo. Given that individual proteins spontaneously phase-separated and produced the telltale inclusion bodies without morphological mixing, inclusion bodies of each cargo protein were biochemically and spatially segregated in the ER lumen. Furthermore, transiently transfected HEK293 cells demonstrated sufficient biosynthetic capacity and membrane plasticity to house four different types of inclusion bodies simultaneously in the ER lumen without compromising cell health.

**Figure 3.**
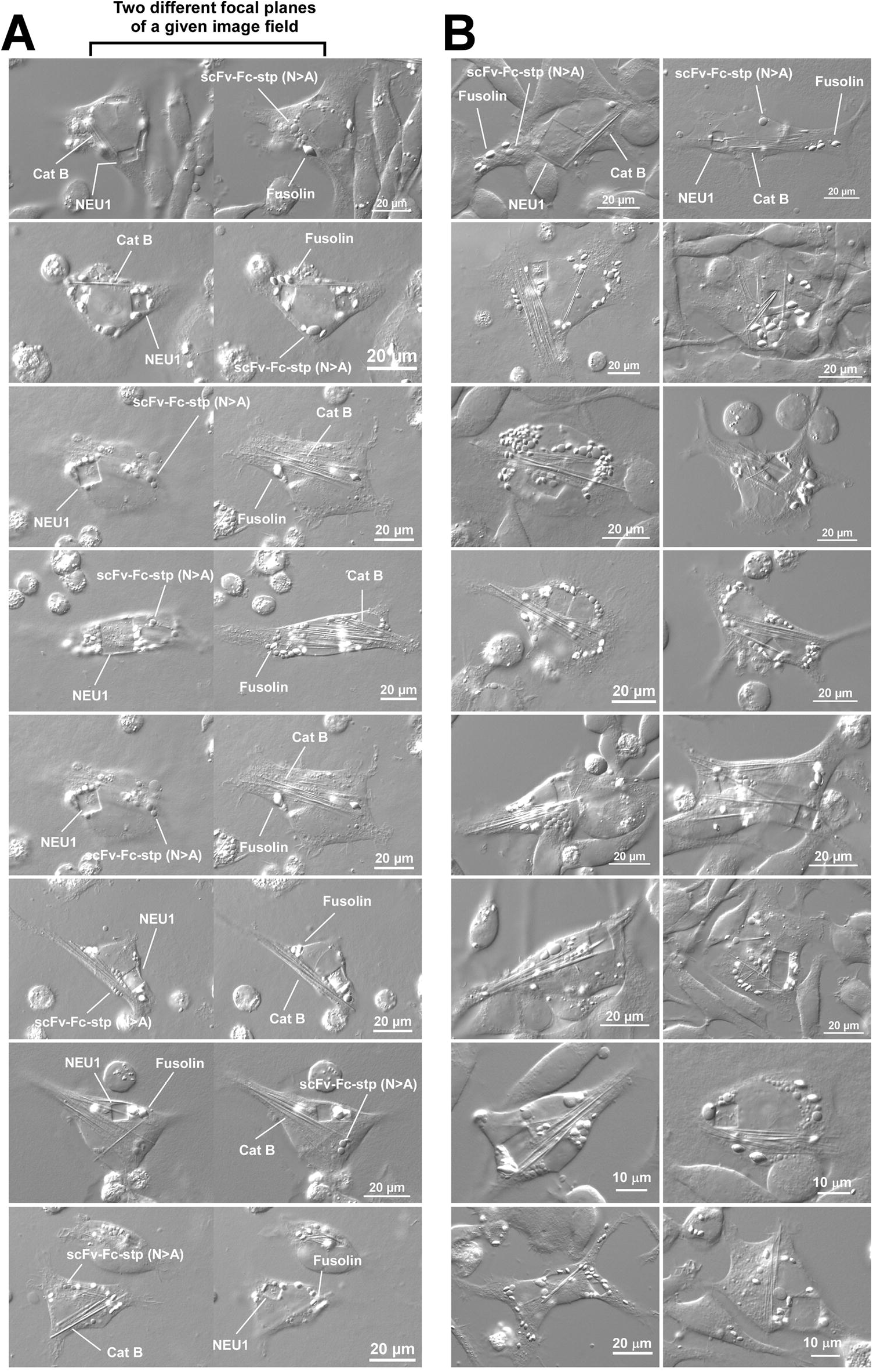
Four separate inclusion bodies can form and co-exist in the ER lumen without morphologically mixed. DIC micrographs of HEK293 cells co-expressing four model cargo proteins simultaneously. On day-4 post-transfection, cells were fixed and imaged. (A) Representative eight image fields captured at two adjacent focal planes are shown side-by-side. Readily distinguishable inclusion body examples are labeled. (B) Sixteen independent image fields showing the cells of interest that house all four inclusion body types in a single focal plane. Representative inclusion bodies are labeled in two images at the top row.

## 3. Discussion

Identifying the four biologically unrelated model cargo proteins made it possible to characterize simultaneously occurring multiple phase separation events in the ER lumen, as well as to explore the limit of a cell’s morphological plasticity and cellular biosynthetic capacity. Co-transfected four cargo proteins independently accumulated in the ER lumen until reaching their respective threshold concentrations and produced characteristic inclusion bodies spontaneously. Because the characteristic inclusion body morphology was preserved during the simultaneous phase separation phenomena, the inclusion body’s identity was readily distinguishable even when all four phase separation events occurred at once. None of the four cargo proteins inhibited the accumulation and phase separation of the other three proteins in the ER lumen, suggesting no dominant □ recessive relationship among the four model cargo proteins. The study additionally showed that intra-ER crystallogenesis and droplet formations do not trigger severe ER stress that otherwise suppresses protein translation mechanisms. Instead, the crystals and droplets continued to grow in size and number over time, a sign that ongoing protein translation steadily supplied the new building blocks and maintained their concentration at saturating levels to support the growth of corresponding inclusion bodies even after the intra-ER inclusion bodies had formed. This observation was clearly different from the accumulation of misfolded/unfolded proteins that induced Russell body-like inclusion bodies by co-aggregating with ER-resident proteins and triggering the unfolded protein responses that suppress cellular protein synthesis [15, 43]. Furthermore, the fact that inclusion body morphology was not mashed up indicated that each cargo protein associates only with its mates to grow the size of respective inclusion bodies in the presence of numerous other proteins in a highly crowded ER lumen. Lastly, the ER membrane is flexible enough and perhaps expandable on demand to accommodate four distinct large inclusion bodies simultaneously without damaging organellar integrity and compromising cell health. As such, transiently transfected HEK293 cells possessed sufficient capacity to safely accommodate at least four distinct inclusion bodies in the ER lumen without acutely disrupting ER homeostasis. The question is whether four different inclusion bodies are the limit of biosynthetic capacity and membrane plasticity. From the standpoint of protein science and material science, it is an interesting topic to explore how many different types of inclusion bodies are producible simultaneously in a single cell or organelle.

In the cases of membrane-less organelle formation by LLPS, functionally antagonizing condensates made of different protein constituents can sometimes coalesce and mix in the cytoplasm to regulate the functions of respective condensates, as shown for the Hippo signaling pathway [44]. By contrast, demixing was the process for membrane-less organelle formation at postsynaptic densities in neurons, where excitatory and inhibitory condensates spontaneously phase-separated into distinct sub-compartments in dendritic spines instead of mixing into one uniform condensate [45]. Yet in another example, the nucleolus is shown to be a multi-phase condensate comprised of three internal liquid phase-separated sub-compartments where different proteins are enriched in each sub-compartment [46]. The four cargo proteins used in this study are biologically and functionally unrelated to each other; thus, there were no known protein-protein interactions or sequence homology among them. They are also from different species or even derived from a synthetic sequence. Because they do not naturally co-exist in the ER lumen of the human HEK293 cells, the results may not directly reflect the biological processes related to health and disease. Nevertheless, these model proteins helped demonstrate that biochemical and spatial demixing underlay the process of simultaneous protein phase separation events in the ER lumen. It is unknown whether the inclusion body demixing widely applies to other sets of phase-separating proteins in the ER lumen. It is also unknown whether mixing or demixing is differentially managed by different sets of phase-separating cargoes or in different subcellular compartments. Furthermore, it is unknown whether it is possible (and if it is, then under what conditions) that two or more distinct inclusion bodies can morphologically mix or merge. In this sense, it is worthwhile designing similar experiments using a set of cargo proteins that (i) have high sequence homology (i.e., disease alleles), (ii) are known to interact with each other, or (iii) function in the same pathway. Such studies may unravel new insights into the underlying mechanisms of heterozygous and multifactorial genetic diseases if wildtype and disease-causing mutants possess different phase-separating propensities.

## 4. Materials and Methods

### 4.1. Detection antibodies and reagents

Rabbit anti-calnexin (cat. C4731) was from Sigma-Aldrich. Rabbit anti-giantin was from Covance (cat. PRB-114P). Mouse anti-LAMP1 (clone H4A3, cat. sc-20011) was from Santa Cruz Biotechnology. All chemical reagents were obtained from Sigma-Aldrich unless specifically mentioned.

### 4.1. Expression constructs

The expression constructs encoding human NEU1 [15], ACEV fusolin [16], and scFv-Fc-stp (N>A) [13] were previously reported. The nucleotide sequence for *Trypanosoma brucei* cathepsin B was generated from the amino acid sequence (UniProt D6XHE1) after optimizing the codon usage to human preference by using a publicly available tool from GENEWIZ (South Plainfield, NJ, USA). The recombinant DNA sequence was verified by the Sanger method and subcloned into a pTT^®^5 mammalian expression vector licensed from the National Research Council of Canada.

### 4.2. Cell culture and transfection

HEK293-6E cell line (herein HEK293) was obtained from the National Research Council of Canada. HEK293 cells were suspension cultured in a humidified CO2 Reach-In incubator (37 °C, 5% CO2) using FreeStyle™ 293 Expression Medium (Thermo Fisher Scientific). The cells were grown in vented cap Corning^®^ Erlenmeyer flasks placed on Innova 2100 shaker platforms (New Brunswick Scientific) rotating at 130 □ 135 rpm. The expression construct was transfected into HEK293 cells using a commonly used polyethylenimine method. Difco yeastolate cell culture supplement (BD Biosciences) was added to the suspension cell culture at 24 hr post-transfection.

### 4.3. Microscopy

Transfected cells in suspension format were seeded onto poly-D-lysine coated glass coverslips at 48 hours post-transfection. Cells were then statically cultured up to day-6 post-transfection in CO2 incubators at 37°C. At a designated time after transfection, cells were fixed in 100 mM sodium phosphate buffer (pH 7.2) containing 4% paraformaldehyde for 30 min at room temperature. Fixed cells were directly used for DIC microscopy or processed for indirect immunofluorescent microscopy. For immunostaining, cells were permeabilized in PBS containing 0.4% saponin, 1% BSA, and 5% fish gelatin for 15 min, followed by incubation with primary antibodies for 60 min. After three washes in the permeabilization buffer, the cells were probed with secondary antibodies for 60 min in the permeabilization buffer. Coverslips were mounted onto slide-glass using Vectashield mounting media (Vector Laboratories) and cured overnight at 4°C. The slides were analyzed on a Nikon Eclipse 80i microscope with a 60× or 100× CFI Plan Apochromat oil objective lens and Chroma FITC-HYQ or Texas Red-HYQ filter. DIC and fluorescent images were acquired using a Cool SNAP HQ2 CCD camera (Photometrics) and Nikon Elements BR imaging software.

## 6. Acknowledgements

HH is personally grateful to Yoko Azumi for her continued support and encouragement.

## References

1. Charcot, J. and C. Robin, Observation de leucocythemie. C. R. Mem. Soc. Biol. (Paris), 1853. 5: p. 44–50.

2. Leyden, E., Zur Kenntniss des Bronchial-Asthma. Virchow’s Arch., Path. Anat., 1872. 54(4): p. 324–344.

3. Reinke, F.B., Beiträge zur histologie des menschen. Archiv für mikroskopische Anatomie, 1896. 47(1): p. 34–44.

4. Glaus, A., Über multiples Myelozytom mit eigenartigen, zum Teil kristallähnlichen Zelleinlagerungen, kombiniert mit Elastolyse und ausgedehnter Amyloidose und Verkalkung. J Virchows Archiv für pathologische Anatomie und Physiologie und für klinische Medizin, 1917. 223(3): p. 301–339.

5. Doye, J.P., W.C.J.C.o.i.c. Poon, and i. science, Protein crystallization in vivo. Current opinion in colloid & interface science, 2006. 11(1): p. 40–46.

6. Schonherr, R., J.M. Rudolph, and L. Redecke, Protein crystallization in living cells. Biol Chem, 2018. 399(7): p. 751–772.

7. Mudogo, C.N., et al., Protein phase separation and determinants of in cell crystallization. Traffic, 2020. 21(2): p. 220–230.

8. Hasegawa, H., Aggregates, crystals, gels, and amyloids: intracellular and extracellular phenotypes at the crossroads of immunoglobulin physicochemical property and cell physiology. Int J Cell Biol, 2013. 2013: p. 604867.

9. Boeynaems, S., et al., Protein Phase Separation: A New Phase in Cell Biology. Trends Cell Biol, 2018. 28(6): p. 420–435.

10. Li, Y., et al., Membraneless organelles in health and disease: exploring the molecular basis, physiological roles and pathological implications. Signal Transduct Target Ther, 2024. 9(1): p. 305.

11. Hasegawa, H., et al., In vivo crystallization of human IgG in the endoplasmic reticulum of engineered Chinese hamster ovary (CHO) cells. J Biol Chem, 2011. 286(22): p. 19917–31.

12. Hasegawa, H., et al., Modulation of in vivo IgG crystallization in the secretory pathway by heavy chain isotype class switching and N-linked glycosylation. Biochim Biophys Acta, 2014. 1843(7): p. 1325–38.

13. Hasegawa, H., N. Patel, and A.C. Lim, Overexpression of cryoglobulin-like single-chain antibody induces morular cell phenotype via liquid-liquid phase separation in the secretory pathway organelles. FEBS J, 2015. 282(15): p. 2777–95.

14. Hasegawa, H., et al., Intermolecular interactions involving an acidic patch on immunoglobulin variable domain and the gamma2 constant region mediate crystalline inclusion body formation in the endoplasmic reticulum. Cell Logist, 2017. 7(3): p. e1361499.

15. Hasegawa, H., Simultaneous induction of distinct protein phase separation events in multiple subcellular compartments of a single cell. Exp Cell Res, 2019. 379(1): p. 92–109.

16. Hasegawa, H., et al., Light chain subunit of a poorly soluble human IgG2lambda crystallizes in physiological pH environment both in cellulo and in vitro. Biochim Biophys Acta Mol Cell Res, 2021. 1868(9): p. 119078.

17. Hasegawa, H., Temperature-dependent intracellular crystallization of firefly luciferase in mammalian cells is suppressed by D-luciferin and stabilizing inhibitors. Exp Cell Res, 2024. 440(1): p. 114131.

18. Lemaire, K., et al., Insulin crystallization depends on zinc transporter ZnT8 expression, but is not required for normal glucose homeostasis in mice. Proc Natl Acad Sci U S A, 2009. 106(35): p. 14872–7.

19. Soragni, A., et al., Toxicity of eosinophil MBP is repressed by intracellular crystallization and promoted by extracellular aggregation. Mol Cell, 2015. 57(6): p. 1011–1021.

20. Karasaki, S., Studies on amphibian yolk 1. The ultrastructure of the yolk platelet. J Cell Biol, 1963. 18(1): p. 135–51.

21. Jiang, L., et al., Biogenesis of the protein storage vacuole crystalloid. J Cell Biol, 2000. 150(4): p. 755–70.

22. Schnepf, E., et al., Bacillus thuringiensis and its pesticidal crystal proteins. Microbiol Mol Biol Rev, 1998. 62(3): p. 775–806.

23. Veenhuis, M., et al., Substructure of crystalline peroxisomes in methanol-grown Hansenula polymorpha: evidence for an in vivo crystal of alcohol oxidase. Mol Cell Biol, 1981. 1(10): p. 949–57.

24. Tsukada, H., Y. Mochizuki, and S. Fujiwara, The nucleoids of rat liver cell microbodies. Fine structure and enzymes. J Cell Biol, 1966. 28(3): p. 449–60.

25. Tenberge, K., et al., Purification and immuno-electron microscopical characterization of crystalline inclusions from plant peroxisomes. Protoplasma, 1997. 196: p. 142–154.

26. Dogan, S., L. Barnes, and W.P. Cruz-Vetrano, Crystal-storing histiocytosis: report of a case, review of the literature (80 cases) and a proposed classification. Head Neck Pathol, 2012. 6(1): p. 111–20.

27. Kmoch, S., et al., Link between a novel human gammaD-crystallin allele and a unique cataract phenotype explained by protein crystallography. Hum Mol Genet, 2000. 9(12): p. 1779–86.

28. Hunt, J.A. and V.M. Ingram, Allelomorphism and the chemical differences of the human haemoglobins A, S and C. Nature, 1958. 181(4615): p. 1062–3.

29. Persson, E.K., et al., Protein crystallization promotes type 2 immunity and is reversible by antibody treatment. Science, 2019. 364(6442): p. eaaw4295.

30. Zhao, Y., et al., Crystalline State Determines the Potency of Galectin-10 Protein Assembly to Induce Inflammation. Nano Lett, 2022. 22(6): p. 2350–2357.

31. Farrants, G.W., S. Hovmoller, and A.M. Stadhouders, Two types of mitochondrial crystals in diseased human skeletal muscle fibers. Muscle Nerve, 1988. 11(1): p. 45–55.

32. Gupta, S.K., et al., Intranuclear Reinke’s crystals in a testicular Leydig cell tumor diagnosed by aspiration cytology. A case report. Acta Cytol, 1994. 38(2): p. 252–6.

33. Shin, M., et al., Drosophila immune cells transport oxygen through PPO2 protein phase transition. Nature, 2024. 631(8020): p. 350–359.

34. Yuan, P., et al., A HEX-1 crystal lattice required for Woronin body function in Neurospora crassa. Nat Struct Biol, 2003. 10(4): p. 264–70.

35. Jedd, G. and N.H. Chua, A new self-assembled peroxisomal vesicle required for efficient resealing of the plasma membrane. Nat Cell Biol, 2000. 2(4): p. 226–31.

36. Adoutte, A., N.G. de Loubresse, and J. Beisson, Proteolytic cleavage and maturation of the crystalline secretion products of Paramecium. J Mol Biol, 1984. 180(4): p. 1065–81.

37. Doye, J.P., A.A. Louis, and M. Vendruscolo, Inhibition of protein crystallization by evolutionary negative design. Phys Biol, 2004. 1(1-2): p. P9–13.

38. Bintrim, S.B. and J.C. Ensign, Insertional inactivation of genes encoding the crystalline inclusion proteins of Photorhabdus luminescens results in mutants with pleiotropic phenotypes. J Bacteriol, 1998. 180(5): p. 1261–9.

39. Bonten, E., et al., Characterization of human lysosomal neuraminidase defines the molecular basis of the metabolic storage disorder sialidosis. Genes Dev, 1996. 10(24): p. 3156–69.

40. Koopmann, R., et al., In vivo protein crystallization opens new routes in structural biology. Nat Methods, 2012. 9(3): p. 259–62.

41. Redecke, L., et al., Natively inhibited Trypanosoma brucei cathepsin B structure determined by using an X-ray laser. Science, 2013. 339(6116): p. 227–230.

42. Harding, H.P., Y. Zhang, and D. Ron, Protein translation and folding are coupled by an endoplasmic-reticulum-resident kinase. Nature, 1999. 397(6716): p. 271–4.

43. Hasegawa, H., et al., Single amino acid substitution in LC-CDR1 induces Russell body phenotype that attenuates cellular protein synthesis through eIF2alpha phosphorylation and thereby downregulates IgG secretion despite operational secretory pathway traffic. MAbs, 2017. 9(5): p. 854–873.

44. Wang, L., et al., Multiphase coalescence mediates Hippo pathway activation. Cell, 2022. 185(23): p. 4376–4393 e18.

45. Zhu, S., et al., Demixing is a default process for biological condensates formed via phase separation. Science, 2024. 384(6698): p. 920–928.

46. Lafontaine, D.L.J., et al., The nucleolus as a multiphase liquid condensate. Nat Rev Mol Cell Biol, 2021. 22(3): p. 165–182.

